# Evidence of a novel sublineage of *Streptococcus agalactiae* in elephants from zoo populations in Germany

**DOI:** 10.1101/2025.03.17.642359

**Authors:** Linda Fenske, Elita Jauneikaite, Maria Getino, Yu Wan, Alexander Goesmann, Tobias Eisenberg

## Abstract

*Streptococcus agalactiae* research primarily centers on investigating human and bovine infections, although this pathogen also can be carried and cause infections in a wider range of animal species. Moreover, infections with *S. agalactiae* are posing significant health implications and, also, recent studies are highlighting a potential zoonotic risk. Despite the comparatively frequent isolation of *S. agalactiae* from elephants, only a few reports document infections in wild and zoo populations. We performed a comparative genomic analysis of 24 elephant isolates from three different zoos in Germany, to achieve a comprehensive characterization. Elephant isolates showed pronounced phylogenetic divergence from isolates of other host species, while also forming clusters based on their zoo of origin and their genotypes (MLST profiles). Capsular serotypes could not be identified for the majority of the isolates (n=20/24). Several genes, associated exclusively with the elephant host may underlie the pathogen’s capacity to improve its survival and virulence across varied ecological niches. This study not only deepens our understanding of *S. agalactiae* across diverse species and environments but also represents the first whole-genome sequencing characterization of *S. agalactiae* isolates from elephants, helping to expand our knowledge about infections in exotic animals.

## Introduction

*Streptococcus agalactiae*, commonly known as Group B streptococcus (GBS), is a facultative anaerobic, non-motile, chain-forming, Gram-positive, catalase-negative bacterium that can cause infections in a wide range of hosts. As a human pathogen, GBS plays a significant role as the primary cause of neonatal infections, usually transmitted from mother to newborn during childbirth, provoking conditions such as pneumonia, meningitis, or septicemia (Le Doare and Heath, 2013). Though GBS is also known to be carried asymptomatically within the urogenital tract and the gut, GBS can also cause infections in adults, especially the elderly; examples of such infections are: urinary tract infections, diabetic foot infections, osteomyelitis, and more severe conditions like toxic shock syndrome, necrotizing fasciitis, disseminated intravascular coagulopathy, and renal dysfunction (Farley and Strasbaugh, 2001; Sendi et al., 2008). GBS is also well known as a pathogen in the dairy industry, where it was first reported as causative agent for the majority of mastitis cases in 1927 (Ruegg, 2017). The implementation of the five-point hygiene plan in the 1960s, which included measures such as rapid identification and treatment of infected cows, whole-herd antibiotic dry cow therapy, post-milking teat disinfection, culling chronically infected cows, and routine disinfection of milking equipment, significantly reduced prevalence of GBS infections but has not eliminated its relevance in the dairy industry today (Hillerton and Berry, 2003; Kabelitz et al., 2021; Mweu et al., 2012). Although published GBS research predominantly focuses on human and bovine diseases, GBS has also been reported to cause infections in a variety of animal species including dolphins (Evans et al., 2006), camels (Seligsohn, 2021), rats (Shuster et al., 2013), fish, seals and other aquatic species (Delannoy et al., 2013; Numberger et al., 2021), and elephants (Eisenberg et al., 2017) .

One of the key virulence factors in GBS is the capsular polysaccharide, which is involved in helping GBS to evade host immune response (Avci and Kasper, 2010). Based on the antigenic properties of the polysaccharide capsule, a serotyping system for GBS was developed (Wilkinson and Moody, 1969). Currently, ten major serotypes (Ia, Ib, II-IX) are recognised, encoded by the capsular locus (*cps*), which comprises 16 to 18 genes (Cieslewicz et al., 2005; Kapatai et al., 2017). It is known that prevalence and distribution of serotypes vary across geographic regions, host species, and clinical presentations (Bianchi-Jassir et al., 2020) and some of the serotypes are also associated with different virulence potential (Burcham et al., 2019; Liu and Liu, 2022). Most studies on GBS focus on human isolates, and though studies in animals have been published, there is a lack of detailed genomic description of GBS isolated from elephants. The first GBS report from elephants dates to 1997 when GBS were identified in pododermatitis lesions from African elephants (Keet et al., 1997) and, then, in Asian elephants in 2008 (Aupperle et al., 2008). Since then, only one other study reported GBS isolated from African and Asian elephants in zoos in Germany (Eisenberg et al., 2017). The latter study employed molecular techniques to characterize and compare the identified GBS isolates from elephants to those in other animals and humans, highlighting the potential genetic diversity of the GBS causing infections in elephants, including the main finding that most of the GBS from elephants were non-typeable using the standard capsular locus typing with multiplex PCR. The non-typable state of GBS isolates is of interest as disease-causing GBS are typically encapsulated, though some studies have reported isolates from humans with absent capsular locus (Creti et al., 2012) and GBS isolates from animals have been reported as non-typable (Sørensen et al., 2019) or not clearly typeable due to previously unknown serotype variants. This could be due to the multiplex PCR designed for human-specific GBS serotypes that does not account for the potential capsular loci diversity of GBS from animals (Crestani et al., 2022).

In the present study, we conducted a comparative genomic analysis of 24 GBS isolates obtained from Asian and African elephants between 2010 and 2023 from zoos in Germany. We aimed to characterize in detail the genomes of these GBS isolates by determining their virulence factors, genotypes based on multilocus sequence typing, capsular serotypes and potential host specific genes. This provides valuable insights into evolution and pathogenicity of GBS populations in less studied host species such as elephants. To our knowledge, this is the first study characterizing GBS isolates from elephants using whole-genome sequencing (WGS).

## Materials and Methods

### Bacterial isolates

A total of 24 GBS isolates from African and Asian elephants in German zoos were whole-genome sequenced and analyzed. Of these, 23 isolates were obtained during routine bacteriological investigations from elephants from two different zoos (A+B) between 2010-2016. Twelve of these isolates were obtained from the previous study by Eisenberg et al., indicated in Supplementary Table 1. One isolate (IHIT53690) was obtained from an elephant in Zoo C in 2023 as part of a routine investigation carried out by the Institute of Hygiene and Infectious Diseases of Animals, Giessen. Metadata of all draft genomes used in this study is provided in Supplementary Table 1.

### Whole-genome sequencing, raw reads processing and assembly

GBS isolates were streaked on Columbia blood agar plates (Oxoid, Basingstoke, UK) and incubated at 37 °C, 5% CO_2_ overnight. Genomic DNA was extracted using the GenElute bacterial Genomic DNA kit (Sigma-Aldrich, Burlington, MA, USA) following the manufacturer’s instructions for Gram-positive bacteria with modifications as follows: GBS was lyzed in 180 μL G+ lysis solution with added 20 μL mutanolysin (Sigma-Aldrich, US; prepared at 3000 U/mL) and 20 μL lysozyme (Sigma-Aldrich, USA; prepared at 100 mg/mL) prior to incubation at 37 °C for 1 h. Subsequent steps followed the manufacturer’s instructions. DNA was quantified using a NanoDrop spectrophotometer (Thermo, Waltham, MA, USA).

For short-read sequencing, multiplexed DNA library preparation was conducted according to the Illumina protocol and WGS was performed on a HiSeq X Ten system (Illumina, USA) with 150-cycle paired-end mode. Raw reads were trimmed and filtered with fastp (v0.23.2) (Chen et al., 2018) and reads were checked for quality with FastQC (v0.11.9) (www.bioinformatics.babraham.ac.uk/projects/fastqc/). Trimmed reads were used for *de novo* assembly into contiguous sequences using Unicycler (v0.5.0) (Wick et al., 2017). The draft assemblies were purged from possible errors with Polypolish (v0.5.0) (Wick and Holt, 2022) and POLCA from the MaSuRCA (v4.1.0) (Maryland Super Read Cabog Assembler) genome assembly and analysis toolkit (Zimin et al., 2013).

Two isolates (161002207-5, 161002207-6) were selected for long-read sequencing using an Oxford Nanopore Technologies (ONT, UK) MinION Mk1B device. Genomic DNA libraries were prepared using the Rapid Barcoding Sequencing Kit (SQK-RBK004; ONT, UK) and loaded into an R9.4.1 flow cell (FLO-MIN106; ONT, UK). Basecalling was conducted using Guppy (v6.5.7) (ONT, UK) under its super-accuracy mode. Long reads were checked for quality using Filtlong (v0.2.1) (https://github.com/rrwick/Filtlong) and assembled using Trycycler (v0.5.5) (Wick et al., 2021) followed by short-read polishing using Polypolish and pypolca (v0.3.1) (Bouras et al., 2024; Zimin and Salzberg, 2020) as recommended in Ryan Wick’s guide to bacterial genome assembly (Wick, 2021). Polished assemblies were annotated using Bakta (v1.7.0) (Schwengers et al., 2021). Contamination and completeness of all genome assemblies were estimated with CheckM2 (v1.0.1) (Chklovski et al., 2023) and furthermore, a taxonomic verification with GTDB-Tk (v2.2.3) was conducted (Chaumeil et al., 2022).

### Population analysis

Multilocus sequence types (MLST) were defined using mlst (v2.23.0) (https://github.com/tseemann/mlst) utilizing the PubMLST *S. agalactiae* database (Jolley et al., 2018); https://pubmlst.org/organisms/streptococcus-agalactiae). Comparative genome analyses for determination of the pan and core genome as well as singleton genes were performed with EDGAR (v3.2) (Dieckmann et al., 2021). For visualization of the core genome phylogeny iTOL (v7) was used (Letunic and Bork, 2021). For phylogenetic placement of the GBS isolates from elephants, a subset of GBS genomes from different host species was selected and included for comparison. Up to 23 representative genomes for each of the different host species were selected from a database of confirmed GBS genomes (Rothen et al., 2019). These included genomes from rats (n=5), dogs (n=4), dolphin (n=1), fish (n=23), frogs (n=2), seals (n=4), camels (n=16), bovines (n=17), and humans (n=23) (Supplementary Table 1). To validate the findings of the phylogenetic analysis, a target-free split k-mer analysis and single-linkage clustering were conducted using SKA (v1.0) (Harris, 2018). Specifically, *k*-mer files (*k*=15) were generated with the fasta subcommand under default parameters; pairwise SNP distances were calculated, and SKA clusters were defined at a threshold of 10 SNPs, provided they satisfied the minimum identity cutoff of 0.9.

### Antimicrobial resistance genes and mobile genetic elements

Antimicrobial resistance genes were detected using AMRFinderPlus (Feldgarden et al., 2019), the Resistance Gene Identifier (RGI) software (Alcock et al., 2023) and abriTAMR (https://github.com/MDU-PHL/abritamr). Putative resistance markers to *β*-lactam antibiotics, were analysed using EDGAR. Orthologous genes encoding the penicillin-binding proteins PBP1A, PBP1B, PBP2A, PBP2B, and PBP2X were identified in the draft genomes using EDGAR. The amino acid sequences of these PBP genes were visualized and manually arranged in Jalview based on sequence similarities (Waterhouse et al., 2009) and subsequently compared to the penicillin-susceptible strain 2603V/R (GenBank: NC_004116). For reconstruction and typing of potential plasmids MOB-suite was used (v3.1.8) (Robertson and Nash, 2018) and the PHASTEST web tool was used to identify prophage regions within the draft genomes (Wishart et al., 2023).

### Analysis of the capsular locus genes

A combination of GBS-SBG (Tiruvayipati et al., 2021), srst2 (v0.2.0) (Inouye et al., 2014), seq_typing (v2.3.0) (https://github.com/B-UMMI/seq_typing), and KMA (v1.4.12) (Clausen et al., 2018) were used to try to determine the capsular serotype based on the capsular loci genes present. As the majority of isolates remained untypeable despite utilizing various tools, an in-depth analysis of the *cps* locus was conducted. The *cps* loci of the elephant GBS isolates were extracted with *in silico* PCR utilizing SnapGene (v8.0.2) (https://www.snapgene.com/). The primers used were introduced in a previous study (Crestani et al., 2022). If no binding sites for these primers were detected within the genomes, alternative primers flanking the *cps* locus at more distant regions were used (Creti et al., 2012).

### Genome-wide association studies

A gene-enrichment analysis, focused on GBS genomes from elephants, was performed using scoary (v1.6.16) (Brynildsrud et al., 2016) from the gene presence/absence matrix generated by panaroo (v1.5.0) (Tonkin-Hill et al., 2020). The latter was generated from the gff3-files annotated with Bakta. The host groups were defined as binary phenotypes with a value of 1 assigned to genomes from the elephant host group and 0 to those GBS genomes from all other species.

## Results

### Sequencing statistics

All 24 genomes were classified as *Streptococcus agalactiae* according to the GTDB taxonomy. The estimated completeness of all draft genomes was above 99.99% with a contamination rate lower than 0.07%. The combined lengths of the assembled contigs range from 1,863,022 to 2,041,196 base pairs (bp) with a GC content between 35.4% and 35.5%. To evaluate how genome size and GC content of the elephant isolates used here integrate into the overall context of GBS genomes, we compared the elephant draft genomes to all GBS genomes currently included in the bacterial web repository BakRep (Fenske et al., 2024). The BakRep v1 contains 10,359 genomes classified as *S. agalactiae* (as of October 2024). After filtering available genomes for completeness of >95% and contamination rate <1%, 9,984 genomes remained for comparison. As none of the genomes included in the repository listed the elephant as the host species, no further subdivision of the data was made. The GBS genomes included in BakRep had a genome size ranging from 1 798 114 to 2 667 783 bp (only two genomes were larger than 2.5 Mbp, i.e. 2.53 Mbp, 2.67 Mbp respectively, in the whole collection) with a GC content ranging from 34.9 to 40.3% (Figure 1). This highlights that the GBS draft genomes from elephants have a mean genome size of 1 930 136 (SD ± 51 596.30) that is below the genomes contained in BakRep with a mean genome size of 2 067 641 (SD ± 69 182.26).

**Figure 1:**
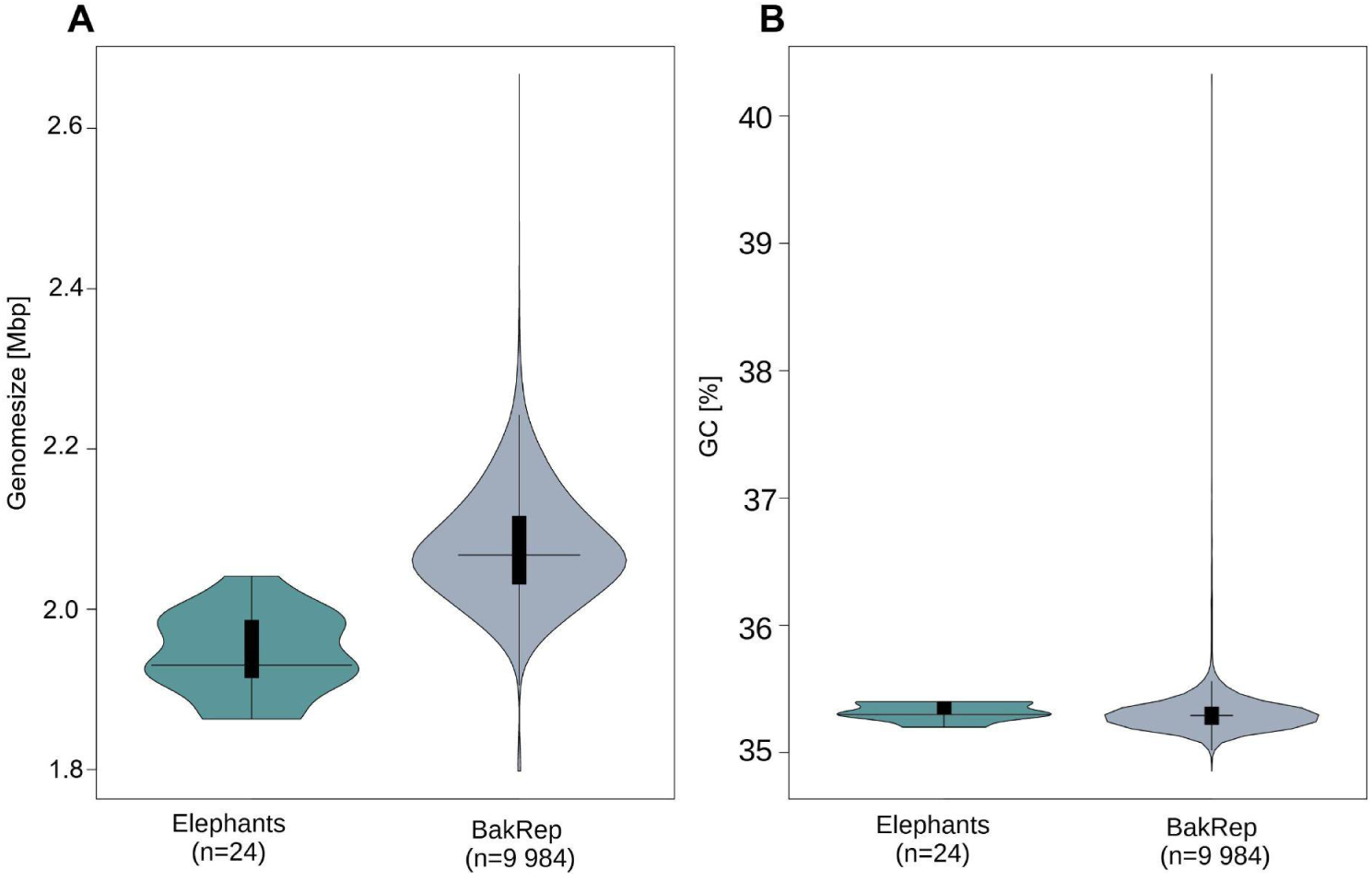
Comparison between genome size (A) and GC content (B). GBS isolates from elephants used in this study (n=24, green) and all GBS genomes included in BakRep (n=9,984, gray). Horizontal black lines in the middle indicate medians; bold black bars represent interquartile ranges; vertical black lines represent outliers.

### Characteristics of GBS genomes from elephants: genotypes, capsular types and AMR

Out of 24 GBS isolates from elephants, 13 (54.17%) were assigned to ST2019 and the remaining 11 isolates were assigned to a new sequence type ST2304 and one isolate was assigned to ST2305, with both STs being first reported in this study and submitted to the *S. agalactiae* MLST database (Table 1). Interestingly, all ST2019 GBS isolates (n=13) came from Zoo A, all ST2304 GBS (n=11) came from Zoo B and one GBS ST2305 isolate came from Zoo C; of note, ST2305 was a single-locus variant (SLV) of ST2304. None of the identified STs could be assigned to a clonal complex.

**Table 1:** MLST results for 24 isolates of *Streptococcus agalactiae* from elephants analyzed in this study utilizing the MLST scheme of Jones *et. al.* (Jones et al., 2003). The newly identified STs in this study are highlighted in bold.

| Isolate | Allele |  |  |  |  |  |  | ST |
| --- | --- | --- | --- | --- | --- | --- | --- | --- |
|  | *adhP* | *pheS* | *atr* | *glnA* | *sdhA* | *glcK* | *tkt* |  |
| 161002207-5 | 16 | 17 | 4 | 2 | 4 | 9 | 5 | **2304** |
| 161002207-6 | 16 | 17 | 4 | 2 | 4 | 9 | 5 | **2304** |
| 161002208-5 | 16 | 17 | 4 | 2 | 4 | 9 | 5 | **2304** |
| 10-7-D-02041 | 20 | 13 | 15 | 1 | 15 | 9 | 5 | 2019 |
| 141000096/2 | 20 | 13 | 15 | 1 | 15 | 9 | 5 | 2019 |
| 141014875 | 20 | 13 | 15 | 1 | 15 | 9 | 5 | 2019 |
| 151001913-2 | 20 | 13 | 15 | 1 | 15 | 9 | 5 | 2019 |
| 151002450-1 | 20 | 13 | 15 | 1 | 15 | 9 | 5 | 2019 |
| 151002450-2 | 20 | 13 | 15 | 1 | 15 | 9 | 5 | 2019 |
| 151003337-1 | 20 | 13 | 15 | 1 | 15 | 9 | 5 | 2019 |
| 151003337-2 | 20 | 13 | 15 | 1 | 15 | 9 | 5 | 2019 |
| 151003441-1 | 20 | 13 | 15 | 1 | 15 | 9 | 5 | 2019 |
| 151003441-2 | 20 | 13 | 15 | 1 | 15 | 9 | 5 | 2019 |
| 151003441-3 | 20 | 13 | 15 | 1 | 15 | 9 | 5 | 2019 |
| 151004802 | 16 | 17 | 4 | 2 | 4 | 9 | 5 | **2304** |
| 151005628-1 | 16 | 17 | 4 | 2 | 4 | 9 | 5 | **2304** |
| 151005628-2 | 16 | 17 | 4 | 2 | 4 | 9 | 5 | **2304** |
| 151006965-1 | 16 | 17 | 4 | 2 | 4 | 9 | 5 | **2304** |
| 151006965-2 | 16 | 17 | 4 | 2 | 4 | 9 | 5 | **2304** |
| 151006967 | 16 | 17 | 4 | 2 | 4 | 9 | 5 | **2304** |
| 151008126 | 20 | 13 | 15 | 1 | 15 | 9 | 5 | 2019 |
| 151010127 | 16 | 17 | 4 | 2 | 4 | 9 | 5 | **2304** |
| 151010216 | 20 | 13 | 15 | 1 | 15 | 9 | 5 | 2019 |
| IHIT53690 | 16 | 23 | 4 | 2 | 4 | 9 | 5 | **2305** |

We knew from a previous study (Eisenberg et al., 2017) that 12 of our GBS isolates from elephants were reported as non-typeable for their capsular serotype using molecular capsular typing methods. The remaining 12 isolates were not routinely serotyped in the laboratory. After WGS, 4 out of 24 (16.7%) GBS isolates were assigned to serotype Ia. For all other isolates no clear assignment could be made even after using various accepted GBS capsular typing methods (Table 2).

**Table 2:**
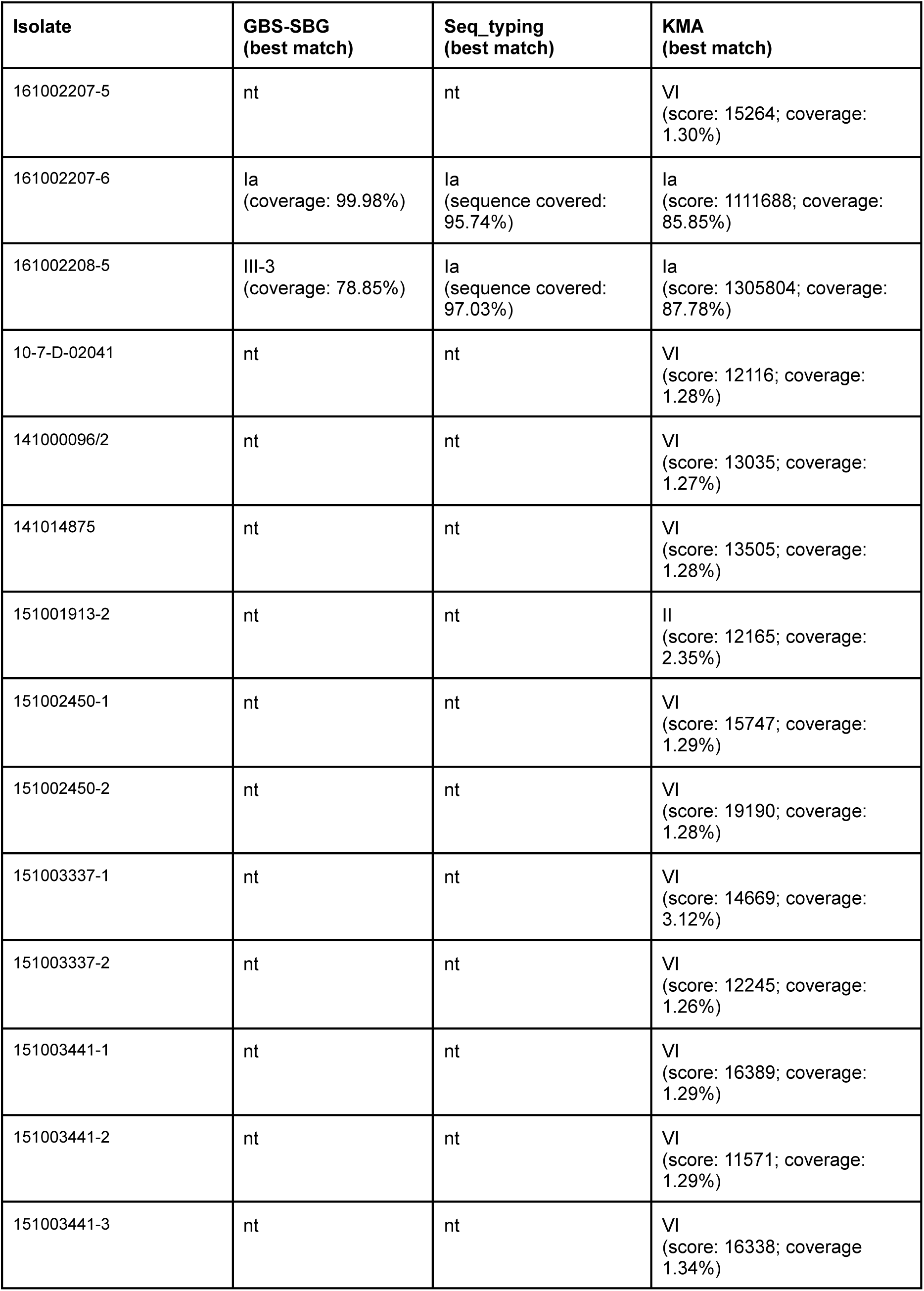
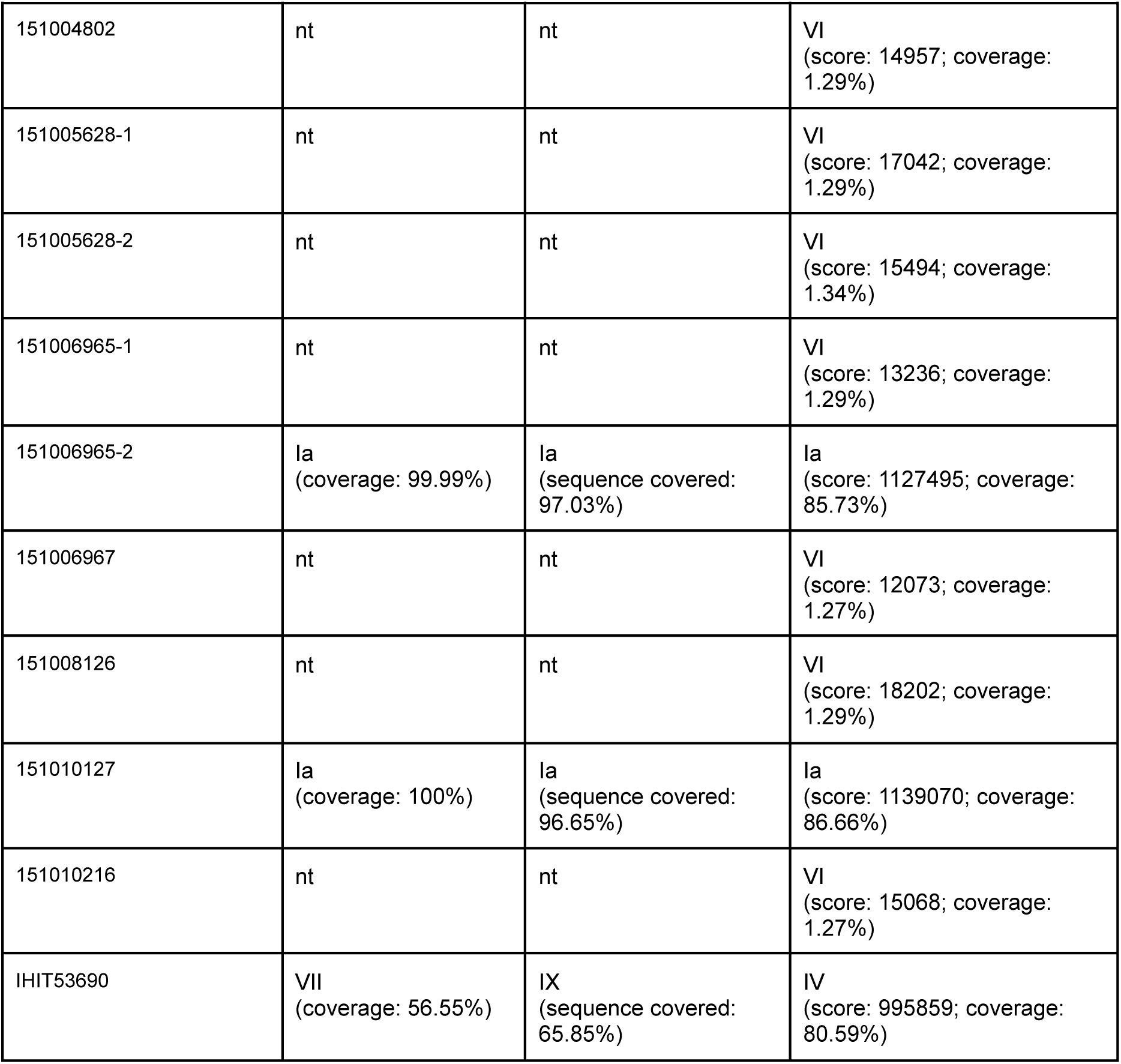
Serotyping results for 24 isolates of *Streptococcus agalactiae* from elephants obtained with different *cps* typing methods.

To analyze the *cps* locus in greater detail *cps* loci of all elephant isolates were extracted. Binding-sites for the primer sequences used by Crestati *et al*. (Crestani et al., 2022) could only be found in the isolates 151006965-2, 151010127, 161002207-6, 161002208-5 and IHIT53690. All isolates, except IHIT53690, were identified as serotype Ia, aligning with the results from other applied methods. The region extracted using primer sequences from Creti *et al*. exhibited a deletion in the remaining 19 isolates, similar to the one previously reported by Creti *et al*. (Creti et al., 2012) (Figure 2). For IHIT53690 no clear serotype assignment could be made due to several gaps and mismatches (Supplementary Figure 2).

**Figure 2:**
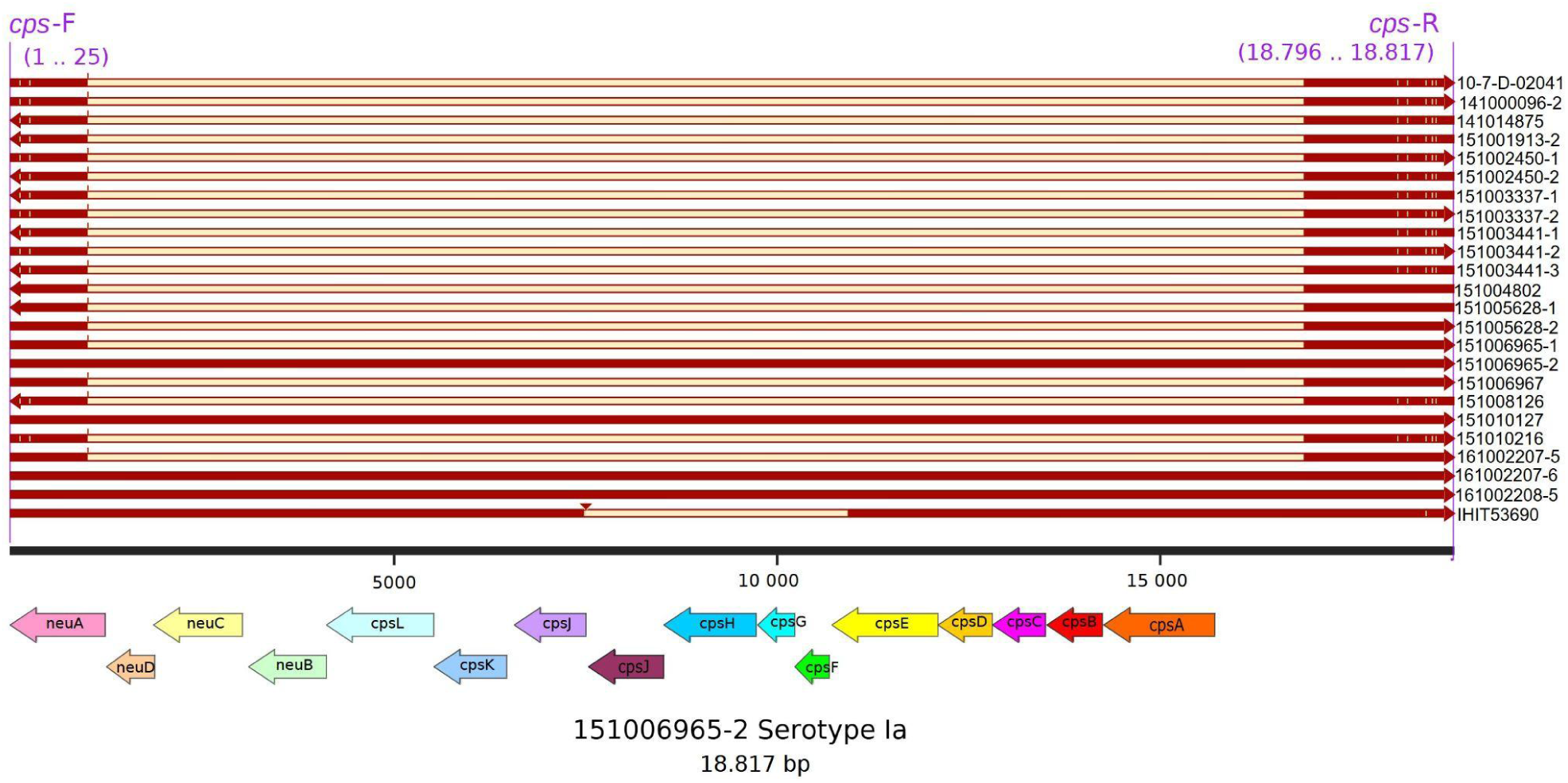
Multiple-sequence alignment of the extracted *cps* locus of the GBS elephant isolates. Colored arrows depict the genetic organisation of the serotype Ia sequence from Isolate 151006965-1, which was used as reference. Positions of primers used are represented at the top. Figure was created with SnapGene and annotation of the genes was performed with Bakta.

Analysis of AMR determinants showed that all 24 GBS isolates were primarily susceptible to antibiotics as the genomes had only the *mprF* gene present, which plays a role in bacterial resistance to cationic antimicrobial peptides (Caliot et al., 2024). In PBP1A all isolates shared a V742A substitution and a deletion of four amino acid residues (739-742) previously described but not associated with reduced susceptibility to *β*-lactams (van der Linden et al., 2020). In PBP1B all isolates had an A95D substitution already detected in GBS from Japan (Li et al., 2020) and in 13/24 (54.17%) isolates a V64I substitution was identified (not previously described). In the PBP2A, 13/24 (54.17%) isolates had the E18K and 10/24 (41.76%) the S394G substitution (not previously described). All isolates showed the V80A substitution in the PBP2B, which was previously detected in other studies (Nagano et al., 2008; van der Linden et al., 2020), while 13/14 (54.17%) isolates had a D572N substitution in the PBP2X gene (not previously described). All substitutions are listed in Supplementary Table 2.

### Mobile genetic elements: prophages identified in GBS isolates

There were neither complete nor partial plasmid sequences identified in any of the 24 isolates. The prophage analysis suggested that 13/24 (54.17%) isolates had predicted intact phages: 7/24 (29.17 %) isolates had *Streptococcus* phage T12 (NC_028700), 2/24 (8.33%) isolates had *Streptococcus* prophage 315.1 (NC_004584.1), 1/24 (4.17%) isolate had intact and one questionable *Staphylococcus* phage SPbeta-like (NC_029119.1), 2/24 (8.33%) isolates had Bacteriophage Shelly (NC_041909.1), 1/24 (4.17%) had *Bacillus* phage G (NC_023719.1), and 1/24 (4.17%) isolate had incomplete *Streptococcus* phage 5093 (NC_012753.1) (Figure 3).

**Figure 3:**
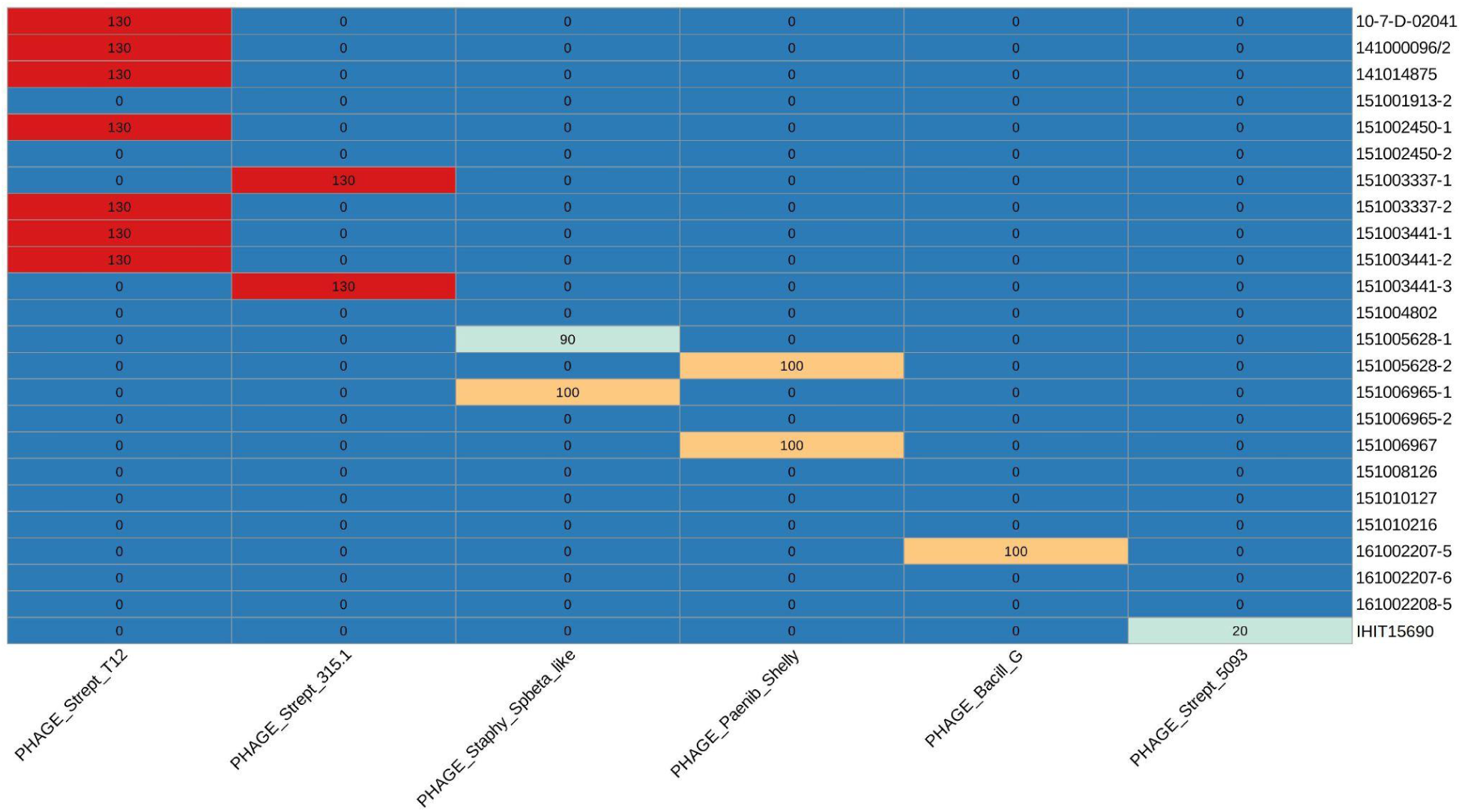
Heatmap of the identified prophage regions. The score (<70: incomplete; 70 - 90: questionable; >90: intact) for each isolate is indicated in the boxes. The heatmap was created with R from the results generated with PHASTEST.

### Pangenome analysis and phylogenetic placement of GBS genomes from elephants

Analysis of the gene content of the 24 elephant GBS isolates revealed a pan genome comprising 2,352 genes, with a core genome of 1,629 genes. When GBS genomes from other host species were included in the analysis, the pangenome expanded to 5,485 genes, while the core genome decreased to 1,325 genes. Within the phylogenetic analysis, the elephant GBS isolates form a distinct sublineage within the same clade as camel isolates but occupy a separate subclade, which also includes another subclade containing two fish isolates and one frog isolate (Figure 4).

**Figure 4:**
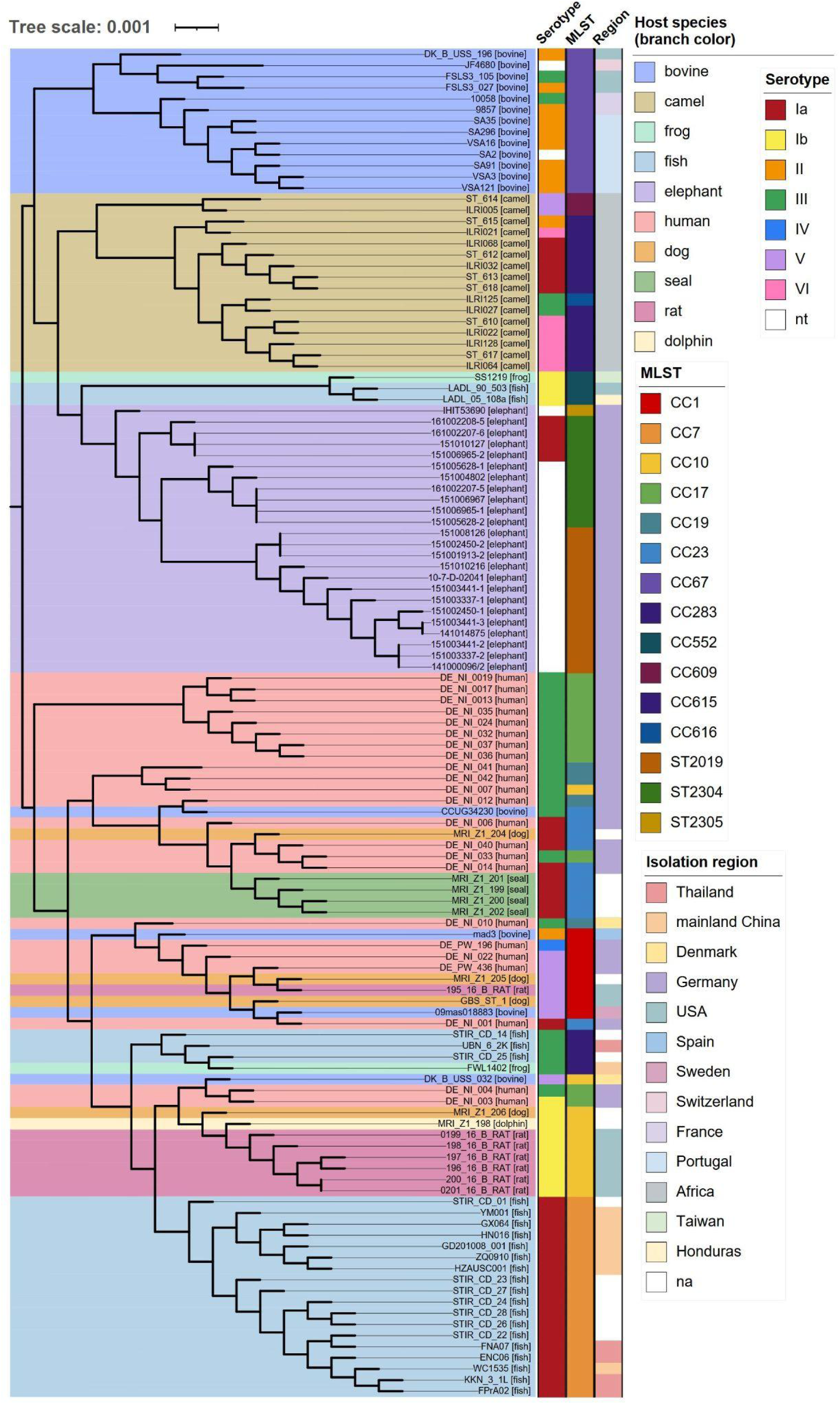
Phylogenetic placement of different GBS isolates (n=121). Tree is color-coded according to the respective host species, with the host species also indicated in square brackets. For creation of the tree the core genes of all genomes were computed with EDGAR. In the following step, alignments of each core gene set are generated using MUSCLE, and the alignments are concatenated to one huge alignment. The tree was constructed with FastTree using the approximately-maximum-likelihood method. The scale bar corresponds to the number of amino acid substitutions per site, with branch lengths reflecting the relative genetic divergence among genomes. Annotation of the tree was done using iTOL.

To further explore the genetic relationships among GBS isolates from elephants, a phylogenetic analysis was conducted exclusively on the 24 elephant GBS isolates (Figure 5). The isolates generally clustered by geographical origin, corresponding to the zoo, from which the host was sampled. An exception was the most recent isolate from an African elephant in Zoo C (collected in 2023), which shared a common ancestor with four GBS isolates from Asian elephants in Zoo A. Notably, a clear clustering pattern emerged based on both the zoo of origin and the collection year. In Zoo B, all isolates were from African elephants and shared the same sequence type (ST2019). These isolates lacked an identifiable capsular serotype and were primarily isolated in 2014. However, this clade also included two isolates from 2015 and one from 2014 that clustered very closely together, suggesting potential persistence of this GBS strain within the zoo population over multiple years.

**Figure 5:**
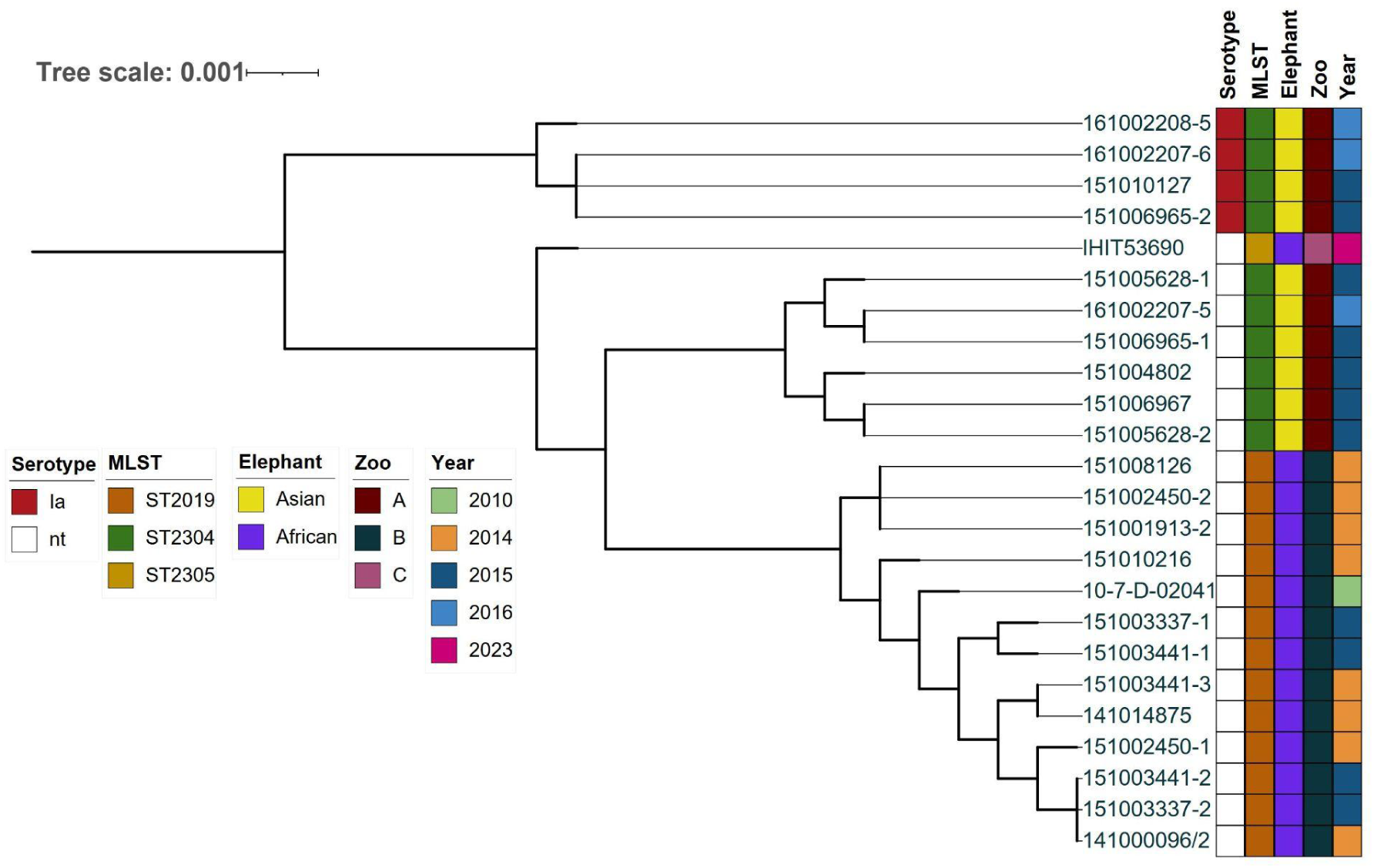
Core genome phylogenetic tree of the 24 elephant isolates. Core genes of all genomes were computed with EDGAR 3.2. based on muscle alignments. Core genes of all genomes are computed with EDGAR. In the following step alignments of each core gene set are generated using MUSCLE, and the alignments are concatenated to one huge alignment. The tree was constructed with FastTree using the approximately-maximum-likelihood method. The scale bar corresponds to the number of amino acid substitutions per site, with branch lengths reflecting the relative genetic divergence among genomes. The tree was annotated using iTol. Colored blocks showing capsular serotype, MLST, elephant species, zoo of origin, isolation year, as indicated by the legend.

With regard to Zoo A, isolates 161002208-5, 151010127, 151006965-2, and 161002207-6 formed a distinct clade separate from the other Zoo A isolates. Notably, these were the only isolates with an identifiable capsular serotype, all classified as serotype Ia. As expected, major clades aligned with MLST types: ST2019 isolates clustered together, while ST2304 clustered with the closely related ST2305 isolate, IHIT53690, as ST2305 is a single-locus variant (SLV) of ST2304 (Figure 5).

### Comparison dataset cluster attribution and genome-wide association investigation

SKA identified a total of 96 clusters among all genomes (n=121), with the GBS isolates from elephants distributed across six distinct clusters (Supplementary Table 3). However, the elephant isolates did not cluster with any isolates from other host species (Supplementary Figure 1). The median SKA SNP distance was 7,147.5 SNPs across all GBS isolates and 2,344.4 SNPs specifically for the elephant isolates. Within Zoo A, the median SNP distance was 182.5 SNPs, while in Zoo B, it was 5.2 SNPs.

A genome-wide association study (GWAS) was conducted using a pan-genome approach to investigate genes that are unique to GBS isolated from elephants. We have found that the *maR* gene was significantly associated with the elephant host, found in all elephant GBS isolates but not in any of the isolates from the other host species. Two additional proteins uniquely associated with all elephant isolates were the Maf family protein and the 3 acyl-CoA thioester hydrolase/BAAT C-terminal domain-containing protein. The Lin0465 protein was found in all but one of the elephant isolates (IHIT53690) and was absent in all but two isolates from other host species (*p*-value: 8.17x10^-15^).

## Discussion

*S. agalactiae* is a known multi-host pathogen, primarily known for causing infections in humans and bovines. Despite this, GBS is also relatively frequently isolated in endangered species such as wild and zoo elephants. However, to date, no studies have investigated these GBS isolates from elephants using WGS. The phylogenetic analysis demonstrates that the core genome of GBS isolates from elephants is distinct from GBS genomes isolated from different host species supporting the niche-specificity of GBS (Crestani et al., 2024; Gori et al., 2020).

There were challenges in defining elephant capsule type, highlighting the overall focus in GBS research on human specific GBS isolates, which are predominantly capsulated and well defined. In our study, serotype Ia was defined only in four GBS isolates with the rest being non-typeable due to a deletion in their capsular locus. A recent study reported a human GBS strain lacking a capsular type, due to the same deletion found in our elephant isolates (Creti et al., 2012). The proposed hypothesis in the mentioned study was that the loss of the capsule occurred due to a recombination event leading to the loss of the whole capsule locus. Given that elephants are a rather uncommon host for GBS and therefore represent a more specialized host niche, capsule expression may not be critical for colonization in this host species. Notably, many GBS vaccines are developed to target specific capsular serotypes, which could allow virulent clones that are not covered by these vaccines to emerge. While these vaccines are intended for human use, this could nevertheless lead to the evolution of vaccine escape mutants. The risk of this happening may increase if there is horizontal transfer of capsular operons between strains or even across different host species (Crestani et al., 2022; Palmeiro et al., 2011).

Eleven of 24 GBS isolates from elephants had a novel ST2304 (n=10) and ST2305 (n=1), highlighting the genetic diversity in the GBS isolated from animals that is still unexplored. Notably, isolate IHIT53690 from Zoo C differed only by one allele from the isolates originating from Zoo B. All other GBS isolates from elephants were classified as ST2019 and were found only in Zoo B. Within the PubMLST database, only another single isolate of ST2019 has been reported, which was also reported as non-typeable for its capsular serotype and has been isolated from elephants in the UK (personal communication, Dr Jauneikaite, manuscript in preparation).

We have not found any acquired antimicrobial resistance genes present in the 24 GBS from the elephants. Penicillin-binding proteins (PBPs), the enzymes essential for the synthesis of peptidoglycan and components of the cell wall in Gram-positive bacteria, have also been investigated in these isolates. Previous reports indicate the specific point mutations in PBPs that are linked to reduced susceptibility to *β*-lactam antibiotics (Piccinelli et al., 2017; van der Linden et al., 2020). We have found a V80A mutation in PBP2B, in all elephant isolates, which has been reported in two studies, although it was not linked to penicillin-non-susceptible GBS (Hu et al., 2018; van der Linden et al., 2020). None of the other substitutions in PBPs identified in this study have been associated with *β*-lactam resistance to date, unfortunately, we do not have phenotypic antimicrobial susceptibility testing done on these isolates to suggest potential effect on the susceptibility to *β*-lactam antibiotics.

Bacteriophages, including prophages, play a crucial role in the evolution of host genomes and are often associated with enhanced infectivity, pathogenicity, and virulence across various bacterial species (Davies et al., 2016; Lichvariková et al., 2020). Seven GBS isolates in our study carried intact prophage regions with most gene hit counts for *Streptococcus* phage T12. This phage is a prototypic temperate phage first associated with *Streptococcus pyogenes,* that carries the *speA* gene coding for erythrogenic toxin A (Weeks and Ferretti, 1984). However, there are no reports of phage T12 in GBS, and none of the elephant isolates in the pan-genome analysis contained the *speA* gene or any related toxin.

When examining the presence of virulence genes, only one gene was consistently found across all elephant GBS isolates in comparison to other GBS isolates. This gene encodes the MprF enzyme, responsible for the unique synthesis of a cationic glycolipid, Lys-Glc-DAG, which aids in the invasion of human endothelial cells, suggesting a potential role in enhancing bacterial entry into host cells and promoting disease progression (Joyce et al., 2022). However, it remains to be confirmed whether it fulfills the same function in elephants. The *maR* gene found exclusively in all elephant isolates encodes the MarR family transcriptional regulator, which plays a key role in controlling the oxidative stress response, a vital environmental sensing function, particularly for pathogenic bacteria (Wilkinson and Grove, 2006). A recent study postulated that the broad host range and ability of GBS to colonize different tissues may be due to the ability of its regulatory systems to recognize and respond to external stimuli, including oxidative and aerobic stress (Wang et al., 2015). Maf-like proteins have been proposed to belong to a family of house-cleaning nucleotide hydrolyzing enzymes, which prevent the incorporation of noncanonical nucleotides into cellular DNA (Galperin et al., 2006). This could also offer an advantage in adapting to different hosts and environmental conditions. Lin0465, which encodes an intracellular PfpI protease in *Listeria*, plays a role in stress responses (Harter et al., 2017; Hein et al., 2011). A study postulated that lin0464, lin0465, and their homologs contribute more to the environmental fitness of these bacteria rather than to their virulence (Hein et al., 2011). In summary, this set of genes unique to the elephant isolates examined here, collectively confers advantages that enhance adaptability to various hosts and environmental conditions. By enabling GBS to respond effectively to diverse physiological and ecological challenges, these genes likely play a role in its ability to adapt to a rare host like the elephant and its survivability in different environments.

## Conclusion

GBS plays an important role in bacterial infections not only in human and bovine diseases, but also in rarer hosts like the elephant. Comparative genomic analysis revealed that, in several aspects, the elephant GBS isolates differ from those of other host species, despite the geographical limitations of our study. To further clarify whether elephants may represent a distinct sublineage of GBS, future studies should include more isolates from elephants across diverse geographic regions. All of this could contribute to a better understanding of the zoonotic potential and pathogenic properties of GBS.

## Acknowledgments

EJ, MG and YW are affiliated with the National Institute for Health Research Health Protection Research Unit (NIHR HPRU) in Healthcare Associated Infections and Antimicrobial Resistance at Imperial College London in partnership with the UK Health Security Agency, in collaboration with, Imperial Healthcare Partners, University of Cambridge and University of Warwick. The views expressed in this publication are those of the author(s) and not necessarily those of the NHS, the National Institute for Health Research, the Department of Health and Social Care or the UK Health Security Agency. YW is a David Price Evans Research Fellow, funded by the Price David Evans endowment to the University of Liverpool. Microbiological work and sample preparation for whole genome sequencing of GBS isolates was undertaken at the Colebrook Laboratory, a facility supported by the NIHR Imperial Biomedical Research Centre (BRC). Part of bioinformatics analysis was performed on equipment purchased as part of MRC CARP fellowship award MR/T005254/1. We would also like to thank Sophie Aurich from the Institute of Hygiene and Infectious Diseases of Animals, Giessen for providing isolate IHIT53690.

## Author contributions

TE, EJ designed and supervised the study. EJ, MG, YW performed the WGS and analyzed the data. LF conducted bioinformatic analysis and interpreted the data. Bioinformatic analyses were supervised by AG. LF wrote the first draft of the manuscript. LF, EJ wrote the next versions of the manuscript draft. AG and EJ acquired funding for the study. All authors critically checked and contributed to the final version of the manuscript.

## Data summary

Raw whole genome sequencing data for elephant isolates used in this study are available in the European Nucleotide Archive (https://www.ebi.ac.uk/ena/browser/home) project accession number PRJEB28328, individual GBS isolate accession numbers have been listed in Supplementary Table 1. Annotations, taxonomic classifications, MLST data, assembly statistics, and associated metadata can also be found in BakRep (https://bakrep.computational.bio/). Reads and assemblies from Oxford Nanopore MinION sequencing for two GBS isolates 161002207-5 and 161002207-6 have been deposited to ENA project accession number PRJEB85776 under accession numbers for raw reads ERR14368735 and ERR14368736 and accession numbers for hybrid assemblies are x1 and y1 respectively.

## Impact Statement

This study provides the first whole-genome characterization of *Streptococcus agalactia*e isolates from elephants, revealing their distinct phylogenetic divergence from GBS isolates of other host species. Our findings suggest that elephant-derived GBS may represent a unique sublineage, with potential adaptations to their host and environment. The absence of identifiable capsular serotypes in most isolates and the presence of host-associated genes highlight the need for further research on the pathogen’s evolution, virulence, and zoonotic potential. Expanding genomic studies to include isolates from broader geographic regions will be crucial for understanding the role of GBS in exotic animals and its potential impact on both wildlife and public health.

---

**Supplementary Figure 1:**
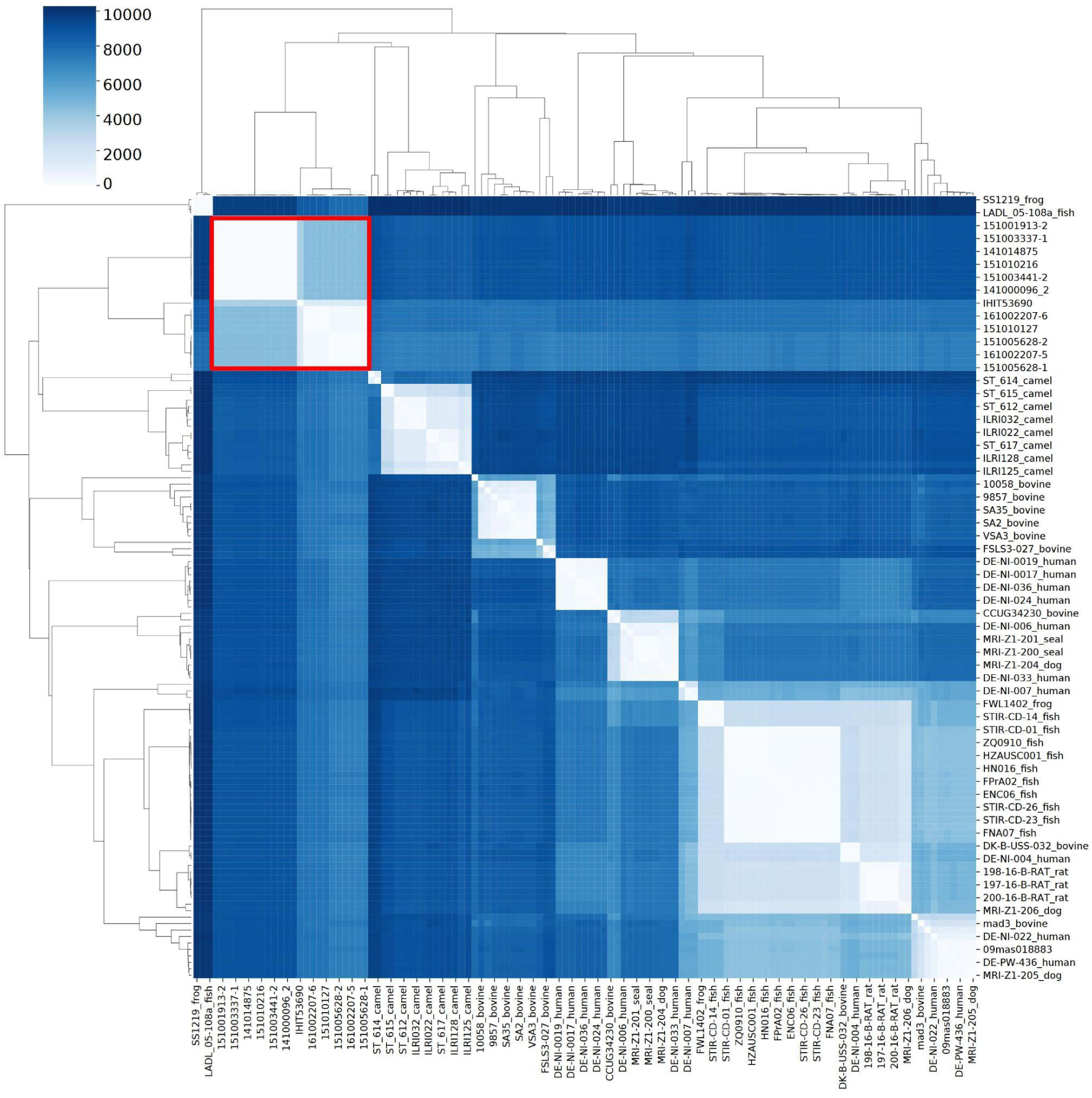
Heatmap with dendrogram displaying the split *k*-mer clustering of various GBS isolates. The distance matrix was created using SKA with a 10 SNP threshold and a minimum identity cutoff of 0.9 and visualized with Python. The elephant cluster is marked with a red square.

**Supplementary Figure 2:**
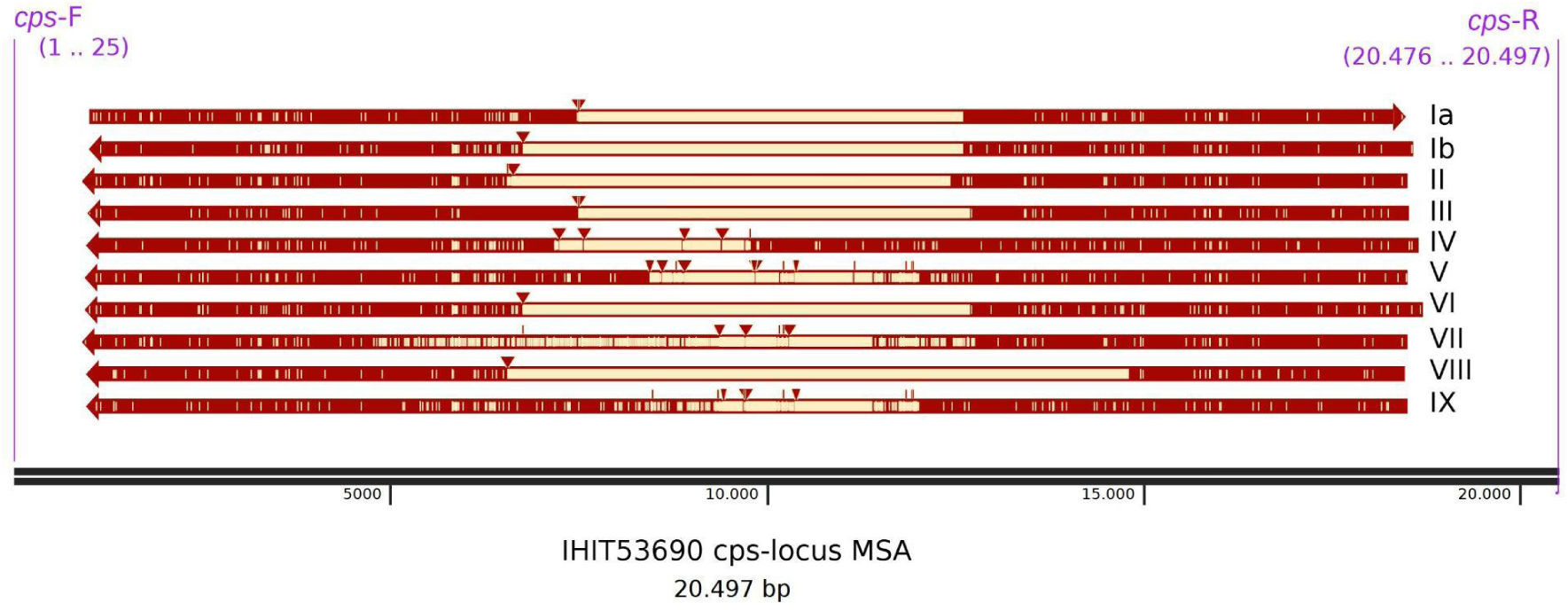
Multiple-sequence alignment of IHIT53690 in comparison with the different *cps* locus variants. Extraction and alignment of the loci was done with SnapGen.

